## Supporting Information Figures and Tables for "Modeling Adsorption, Conformation, and Orientation of the Fis1 Tail Anchor at the Mitochondrial Outer Membrane"

### S.I. SIMULATED ANNEALING

In standard molecular dynamics simulations with Fis1(TA) we could not observe the insertion of the peptide into the membrane within 500 ns. Therefore, we applied simulated annealing (SA), where only the peptide was tempered. Fis1(TA)'s temperature was raised from 298 K to 800 K in 2.5 ns, kept at 800 K for 2.5 ns, cooled down to 298 K in 2.5 ns, followed by an additional 12.5 ns simulation at 298 K. The rest of the system was kept at 298 K during the SA procedure. The annealing cycles were applied for a total simulation time of 1000 ns. Fig. S1 shows the temperature of the peptide (blue), temperature of the system (red) and the potential energy (blue) for the first five annealing cycles.

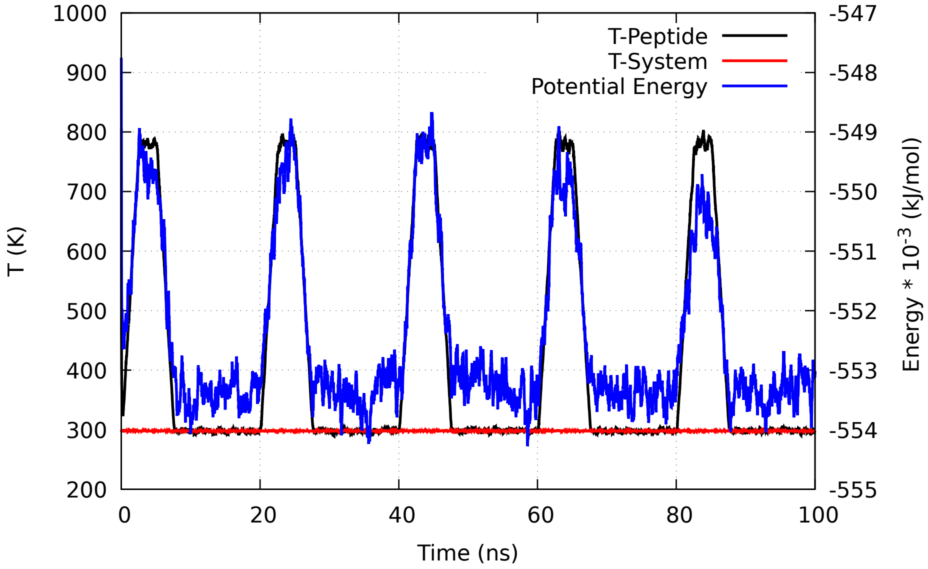


FIG. S1: Temperature of the peptide (T-peptide), temperature of the system (water, lipids and ions; T-System) and potential energy profile during the simulated annealing cycles. Only the first 5 cycles are shown.

Considering the stochastic nature of the insertion process, we have performed an additional set of 10 trials by using the same SA procedure with random seeds. Analysis of all eleven runs in terms of the orthogonal distance from the center-of-mass of the peptide to the membrane centerline and secondary structure evolution during the run are shown in Fig. S2. The data for center-of-mass distances are smoothed by calculating the running average (with a window size of 100 data points).

Nine out of eleven runs display successful insertion of the peptide into the membrane within 1 $\mu s$. Among them eight of them show significant alpha-helix content as revealed by the DSSP analysis. In comparison to the wildtype Fis1(TA), the mutants A144D and L139P show less penetration into the membrane and lower alpha-helix content in general (Fig. S3 and Fig. S4). The extra negative charge in the A144D mutant creates an additional barrier for penetration to the interior of the membrane. In case of L139P, the helix breaker nature of the proline residue destabilizes the secondary structure formation.

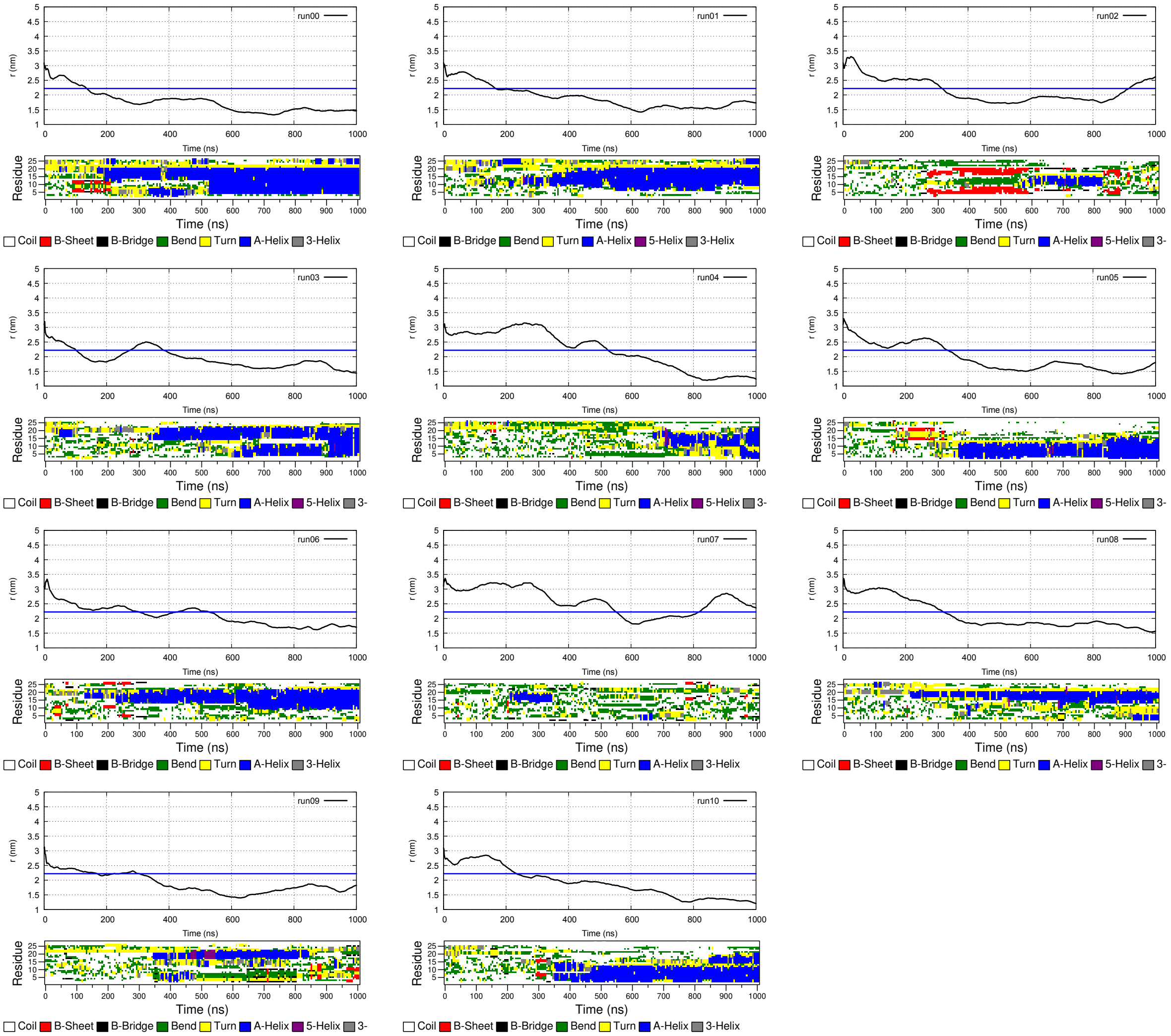


FIG. S2: Analysis of SA runs for wildtype Fis1TA. The results for all eleven trials with different initial random seeds are shown with labels run00-run10. Orthogonal distance from the center-of-mass of the peptide to the membrane center-line (top) and the secondary structural analysis via DSSP (bottom) are shown as a function of the simulation time. The position of the water/membrane interface (defined as the point where the lipid density reaches 500 $kg/m^3$) is marked by the blue solid line.

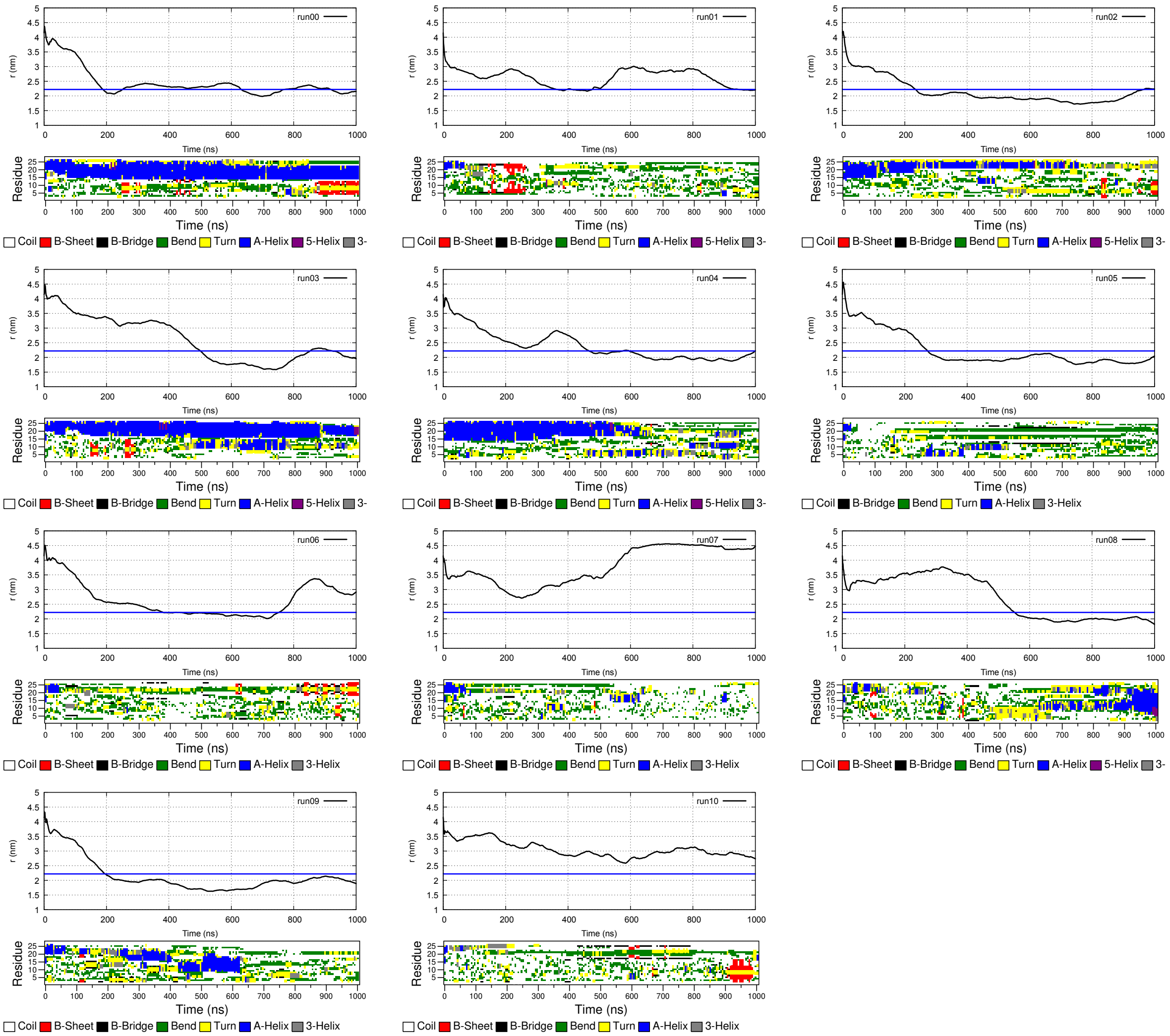

FIG. S3: Analysis of SA runs for the A144D mutant. See Fig. S2 for details.

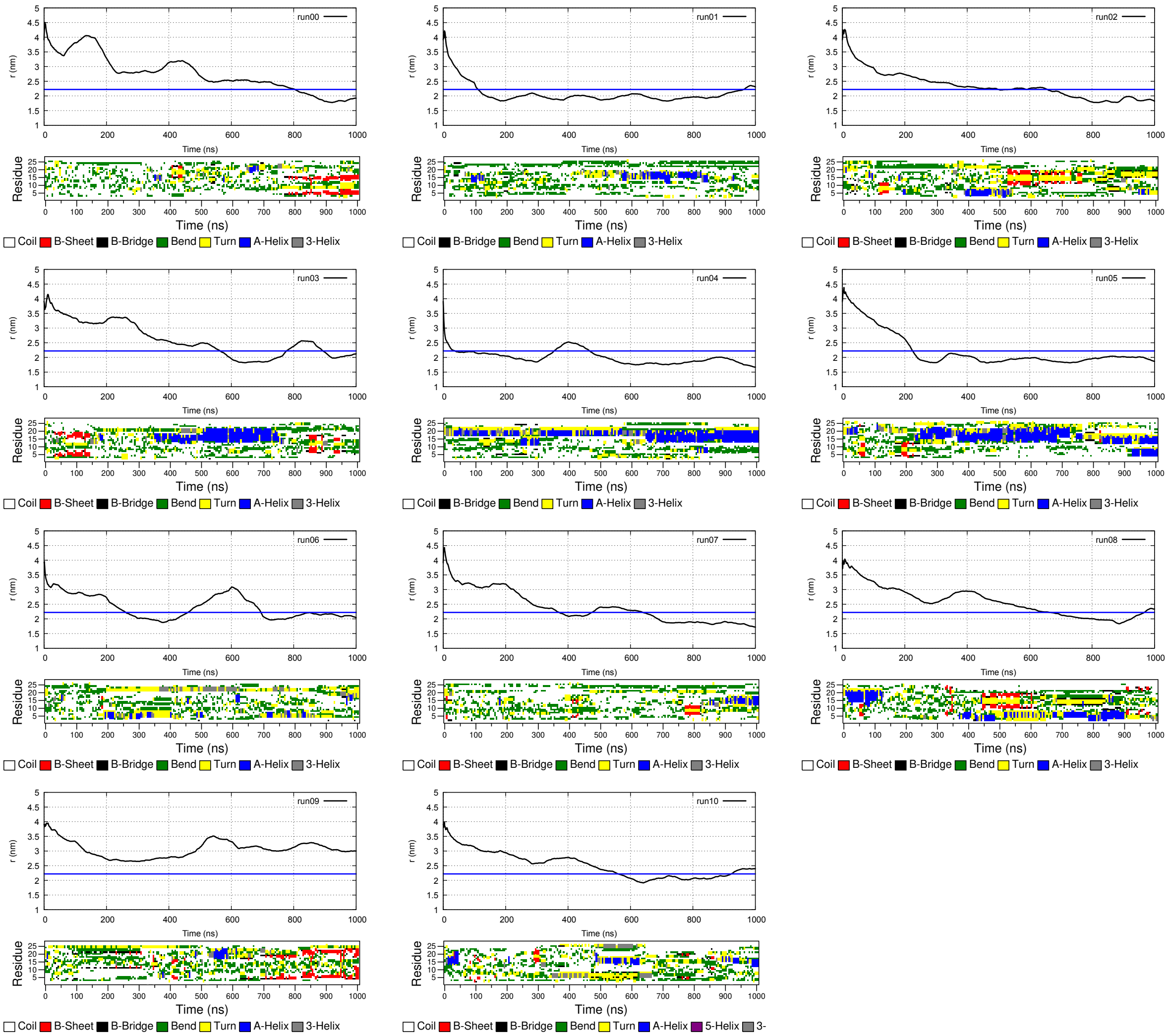

FIG. S4: Analysis of SA runs for the L139P mutant. See Fig. S2 for details.

### S.II. SECONDARY STRUCTURE ANALYSIS FOR THE AA-REX SIMULATION

The atomistic replica exchange molecular dynamics simulations allowed us to analyze the conformational sample space of the peptide within the membrane. We have used 80 replicas spanning a temperature range of 298-471 K for our AA-REX simulation. In Fig. S5 and Fig S6 secondary structure analysis via DSSP are shown for five different temperatures and five different replicas. Even at 471 K the peptide displays a significant alpha-helix propensity. As the replicas go through different temperatures, complete folding and unfolding of the peptide is observed. The one microsecond simulation is sufficient to yield a converged average alpha-helix propensity. The DSSP analysis for selected replicas shown in Fig. S6 demonstrate that the REX simulation is successful in forcing the peptide to alter its conformation as it visits replicas with different temperatures. However, the accessible simulation times are still not sufficient to observe a high number of transitions for each replica.

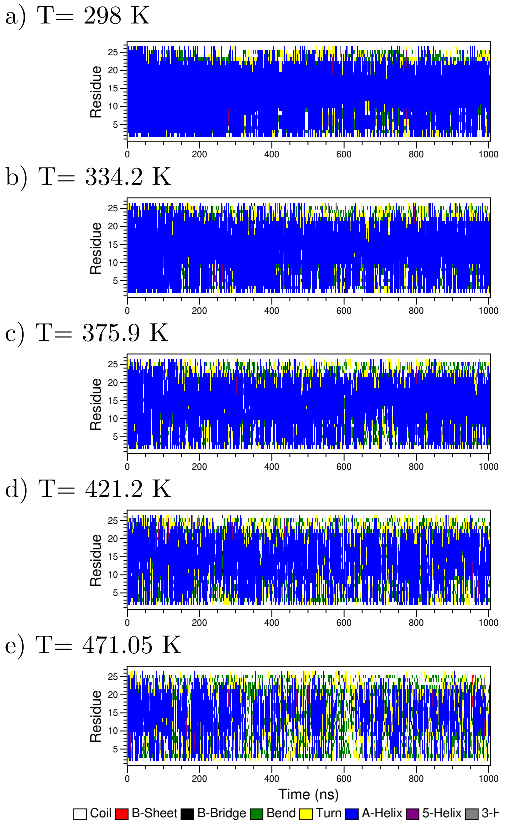


FIG. S5: DSSP analysis of the wild-type Fis1(TA) AA-REX simulation at five different temperatures.

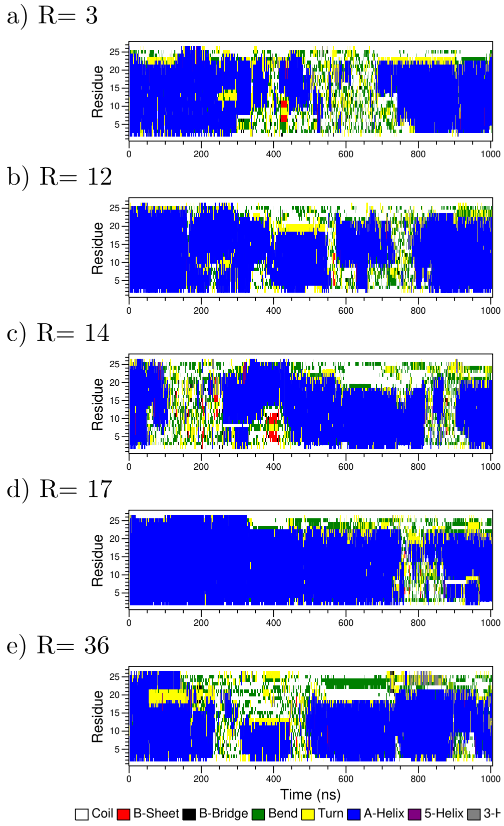


FIG. S6: DSSP analysis of the wild-type Fis1(TA) for five different replicas during the 1 $\mu s$ AA-REX simulation.

### S.III. MARTINI SIMULATION

In order to facilitate the transitons between the monotopic and bitopic configurations of Fis1-TA peptide, we reduced the polar/charged character associated with lysine, arginine and asparagine residues. To this end, we used thirteen replicas (Table I) where the default MARTINI bead types (of both backbone and side chain) of lysine, arginine and asparagine residues are replaced by bead types of less polar (more hydrophobic) character. In the final configuration replica 12 corresponds to the wild-type Fis1 TA peptide and replica 0 represents a TA sequence that is fully hydrophobic in the context of MARTINI bead types.

TABLE I: MARTINI bead types used in each replica of well-tempered metadynamics simulations coupled with Hamiltonian exchanges. BB and SC represent backbone and side chain bead types, respectively. chg-K and chg-R columns demonstrate the charges of lysine and arginine side chain groups. Replica 12 corresponds to the wild-type Fis1 TA peptide. The default MARTINI bead types of polar and/or charged nature for Lysine, Arginine and Asparagine are gradually replaced by uncharged, less polar and/or hydrophobic beads as we move down from replica 12 (wild-type) to replica 0 (the most hydrophobic).

| **Replica id** | **BB-K** | **BB-R** | **BB-N** | **SC-K** | **SC-R** | **SC-N** | **chg-K** | **chg-R** |
|---|---|---|---|---|---|---|---|---|
| 0 | N0 | N0 | N0 | AC1 | AC1 | AC1 | 0.0 | 0.0 |
| 1 | N0 | N0 | N0 | C3 | C3 | C3 | 0.0 | 0.0 |
| 2 | N0 | N0 | N0 | C5 | C5 | C5 | 0.0 | 0.0 |
| 3 | N0 | N0 | N0 | N0 | N0 | N0 | 0.0 | 0.0 |
| 4 | N0 | N0 | N0 | P1 | P1 | P1 | 0.0 | 0.0 |
| 5 | N0 | N0 | N0 | P3 | P3 | P3 | 0.0 | 0.0 |
| 6 | N0 | N0 | N0 | P5 | P5 | P5 | 0.0 | 0.0 |
| 7 | N0 | N0 | N0 | Qd | Qd | P5 | 0.0 | 0.0 |
| 8 | P2 | P2 | P2 | Qd | Qd | P5 | 0.0 | 0.0 |
| 9 | P5 | P5 | P5 | Qd | Qd | P5 | 0.0 | 0.0 |
| 10 | P5 | P5 | P5 | Qd | Qd | P5 | 0.33 | 0.33 |
| 11 | P5 | P5 | P5 | Qd | Qd | P5 | 0.66 | 0.66 |
| 12 | P5 | P5 | P5 | Qd | Qd | P5 | 1.0 | 1.0 |

The convergence check for the well-tempered metadynamics simulation coupled with Hamiltonian replica exchange was done using several metrics. The height of the Gaussian biases added to the system is shown in Fig. S7 for the replica corresponding to the wildtype system. Over time the heights of the added Gaussian peaks decrease as expected from a well-tempered metadynamics simulation. Since we have a two dimensional surface, occasional increases are observed in height.

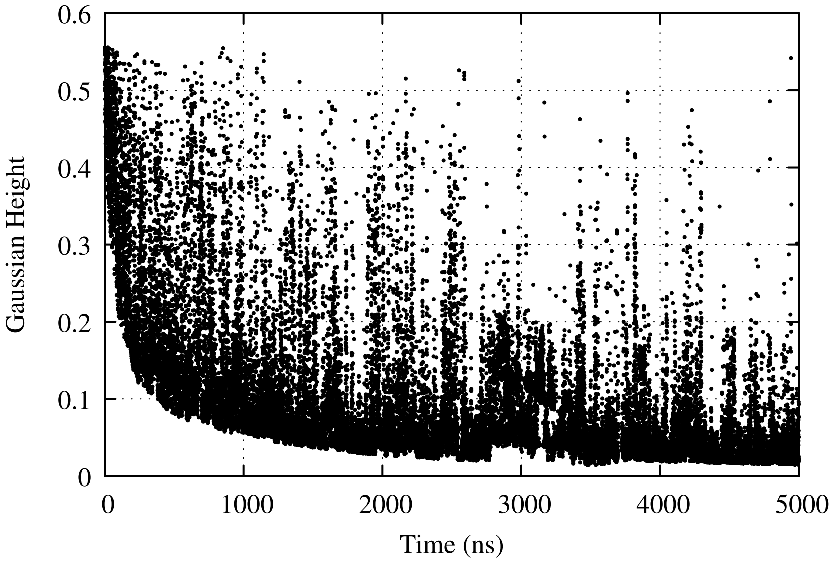


FIG. S7: Height of the Gaussian bias during the 5 $\mu$s well-tempered metadynamics simulation.

Two different collective variables were used for the metadynamics simulation: the orthogonal distance from the peptide center-of-mass to the membrane centerline ($r$) and tilt angle ($\theta$) which measures the angle between the membrane normal (approximated by the z-axis in the simulation box) and the helix axis. In Fig. S8 and Fig. S9 the convergence analysis is shown for the free energy surface projected onto $r$ and $\theta$, respectively. The total simulation time was divided into ten blocks (0.5 $\mu$s each), and evolution of the free energy surface was calculated by using the data from the start till the end of the block. Only the last five blocks are shown in the figures. For both of the collective variables the fluctuations in the free energy profiles between different blocks are less than 2 kJ/mol.

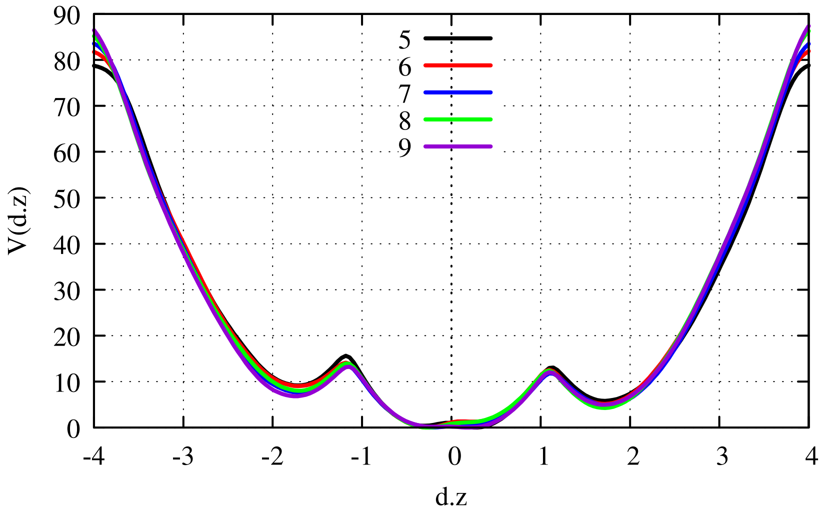


FIG. S8: Convergence of the free energy surface projected onto the collective variable $r$.

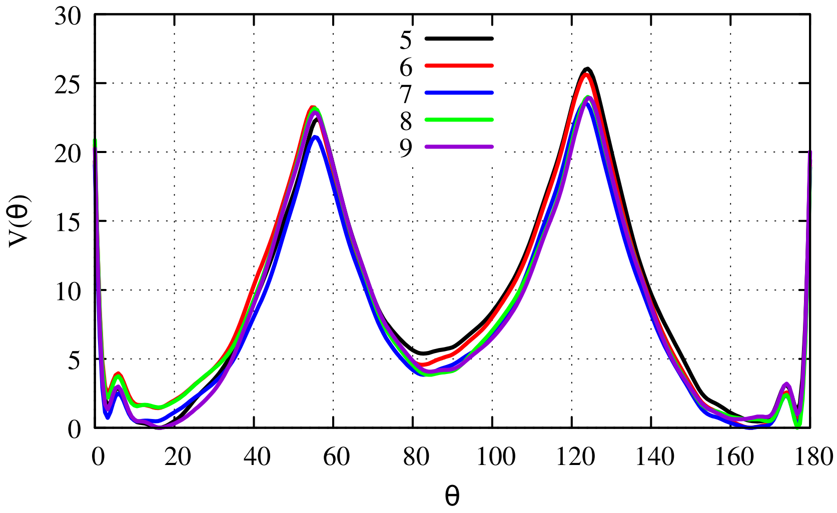


FIG. S9: Convergence for the free energy surface projected onto the collective variable $\theta$.

The evolution of the two dimensional free energy surface for the collective variables $r$ and $\theta$ during the $5\mu s$ simulation is also given in Fig. S10. Similar to the one dimensional analysis the data is divided into ten blocks. In the figure the blocks 0, 3, 6, 7, 8, and 9 are shown for the replica corresponding to the wild-type system.

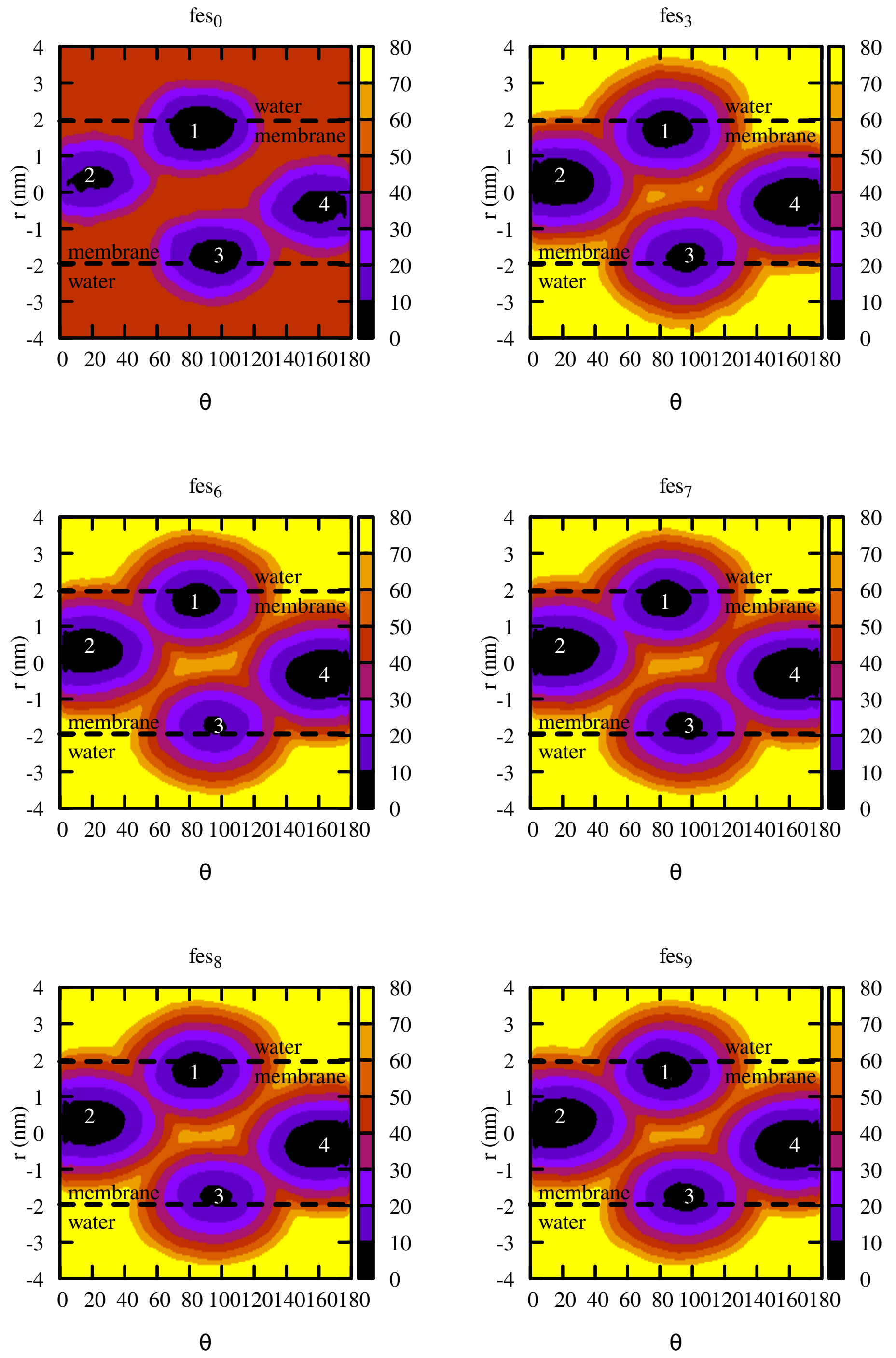


FIG. S10: Analysis of the convergence of the two dimensional free energy surface for the MARTINI simulation. Evolution of the free energy surface for the replica corresponding to the wildtype system is shown.

As explained in the maintext, the wildtype sequence was mutated to a completely hydrophobic sequence to allow the system to explore the conformational space efficiently. In Fig. S11 the final two dimensional free energy surfaces are shown for the replicas 0, 4, 8 and 12. Replica 0 corresponds to the completely hydrophobic mutant and replica 12 corresponds to the wildtype Fis1(TA). Compared to the energy barriers in the wildtype system's free energy surface, the hydrophobic molecule (replica 0) displays significantly lower energy barriers for the transition between monotopic and bitopic orientations. In addition, the difference in barrier height for 1-to-2 and 1-to-4 (similarly 3-to-4 and 3-to-2) pathways is significantly reduced in replica 0.

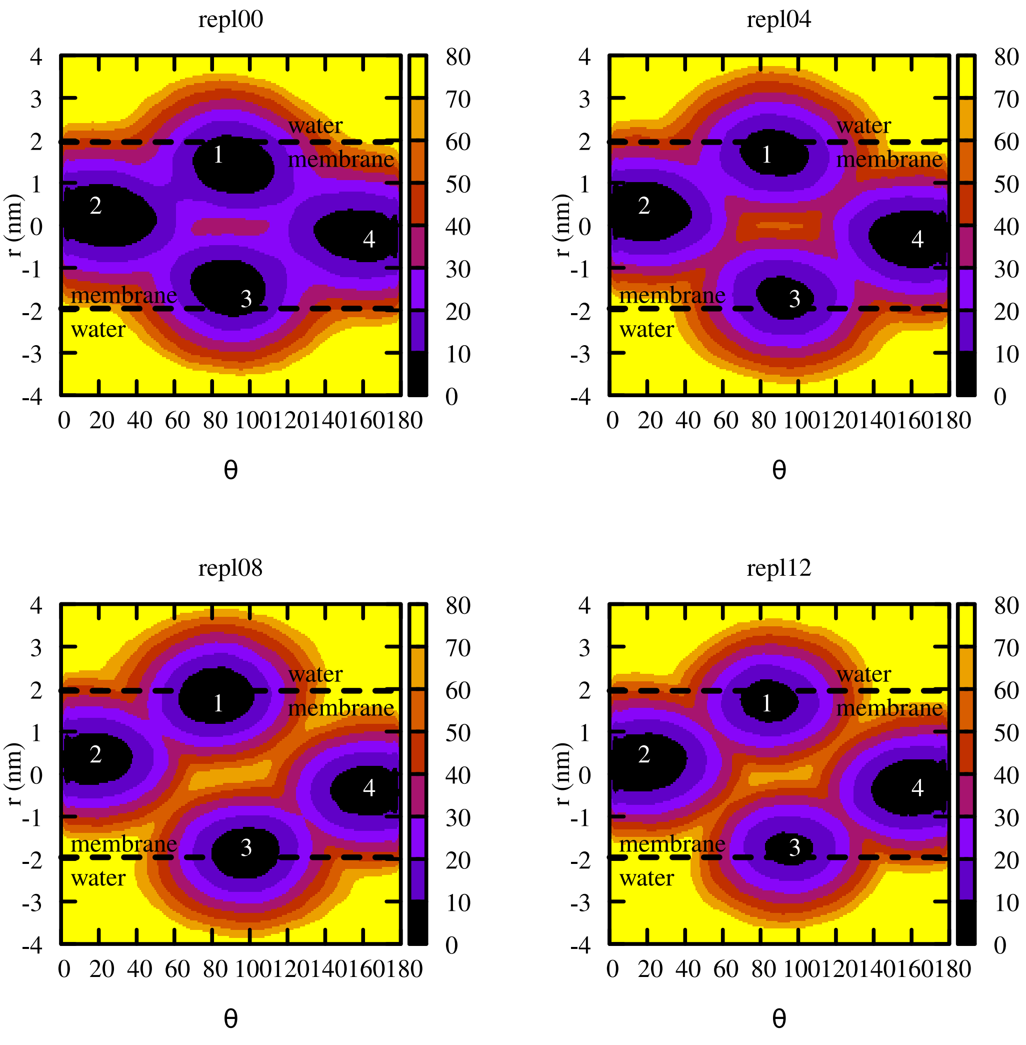


FIG. S11: The final two dimensional free energy surfaces for the replicas 0, 4, 8 and 12. Replica 0 corresponds to the completely hydrophobic mutation and replica 12 corresponds to the wildtype system.
